# Cis-regulatory evolution of the potassium channel gene *kcnj13* during pigment pattern diversification in *Danio* fish

**DOI:** 10.1101/2022.12.05.519077

**Authors:** Marco Podobnik, Ajeet P. Singh, Zhenqiang Fu, Christopher M. Dooley, Hans Georg Frohnhöfer, Magdalena Firlej, Hadeer Elhabashy, Simone Weyand, John R. Weir, Jianguo Lu, Christiane Nüsslein-Volhard, Uwe Irion

## Abstract

Teleost fish of the genus *Danio* are excellent models to study the genetic and cellular bases of pigment pattern variation in vertebrates. The two sister species *Danio rerio* and *Danio aesculapii* show divergent patterns of horizontal stripes and vertical bars that are partly caused by the evolution of the potassium channel gene *kcnj13*. In *D. rerio, kcnj13* is required in melanophores for interactions with xanthophores and iridophores, which cause location-specific pigment cell shapes and thereby influence colour pattern and contrast. Here, we show that cis-regulatory rather than protein coding changes underlie *kcnj13* evolution between the two species. *D. aesculapii* express lower *kcnj13* levels and exhibit low-contrast patterns similar to *D. rerio* mutants. Our results suggest that homotypic and heterotypic interactions between the pigment cells and their shapes diverged between species by quantitative changes in *kcnj13* expression during pigment pattern diversification.

## Introduction

Teleost fish produce some of the most diverse pigment patterns in nature, which are of great evolutionary importance as direct targets of natural and sexual selection. Closely related species of the genus *Danio*, including the widely used model organism zebrafish, *Danio rerio*, develop amazingly different patterns and are therefore excellent models to investigate the evolution of pigment pattern diversification in vertebrates^1-5^. Recently, the phylogenetic relationships in the *Danio* genus have been resolved, which led to the insight that a complex evolutionary history underlies their speciation and morphological diversification^6^.

The horizontally striped pattern in *D. rerio* emerges during metamorphosis when multipotent pigment cell progenitors derived from stem cells located at the dorsal root ganglia (DRGs) migrate into the skin^7-9^. Here they differentiate and form the pattern, presumably by a self-organizing process dependent on multiple cell-cell interactions^10,11^. These interactions lead to the acquisition of location-dependent cell shapes, compact/yellow xanthophores and dense/reflective iridophores in the light stripes, and stellate xanthophores and loose/blue iridophores in the dark stripes^7,12,13^. Melanophores are restricted to the dark stripes. Precise superimposition of the differentially shaped pigment cells is required for colour and contrast of the pattern. The cellular interactions are, at least partially, mediated by direct cell-cell contacts through gap junctions, adhesion molecules and ion channels. Gap junctions are formed by two connexins (Gja4 and Gja5b)^14-16^, Igsf11 and Jam3b regulate adhesion^17,18^ and Kcnj13 is an inwardly rectifying potassium channel^19^. The diverse patterns in other *Danio* fish are produced by the same three types of pigment cells; however, the genetic and cell biological basis of the pattern variation is still largely unexplored. So far, the evolution in two separate cell differentiation pathways, xanthophore-specific Csf1 signalling in *D. albolineatus* and iridophore-specific Edn signalling in *D. nigrofasciatus*, has been linked to patterning differences^20-22^. This mode of evolution might partly cause changes in the timing and strength of the interactions between pigment cells, with cascading effects on their final distribution within the skin.

In this study, we focus on the diversification of pigment patterns between the two sister species *D. rerio* and *D. aesculapii*. Whereas *D. rerio* develop a very stereotypic pattern of sharp horizontal dark and light stripes on the flanks and in the anal and tail fins (Fig. 1a), in *D. aesculapii* a more variable pattern of vertical bars with lower contrast is formed anteriorly on the flank that dissolves into irregular spots posteriorly; the fins are not patterned, except for one dark stripe in the anal fin (Fig. 1b). We have shown that the potassium channel gene *kcnj13* evolved to contribute to these patterning differences between the two species^23^.

**Fig. 1:**
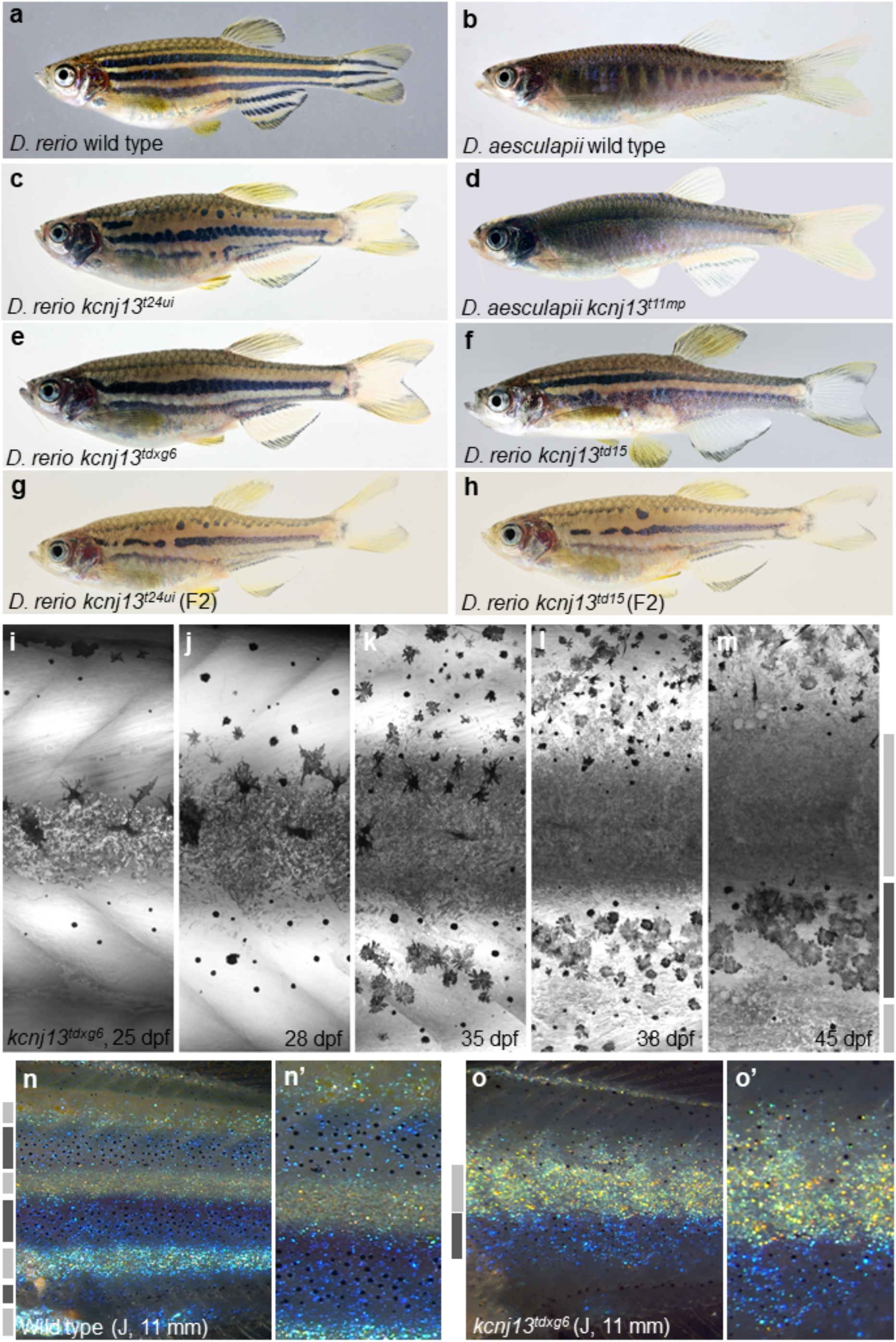
Pigment patterns in wild type and *kcnj13* mutant *D. rerio* and *D. aesculapii*. Pigment patterns in **a** *D. rerio* wild type, **b** *D. aesculapii* wild type, **c** *D. rerio kcnj13*^*t24ui*^, **d** *D. aesculapii kcnj13*^*t11mp*^, **e** *D. rerio kcnj13*^*txg6*^ and **f** *D. rerio kcnj13*^*td15*^. *kcnj13*^*t24ui*^ and *kcnj13*^*td15*^ were crossed to produce trans-heterozygous *kcnj13*^*tui24/td15*^ F1 fish (not shown), which were then incrossed to generate F2 fish with the genotypes **g** *kcnj13*^*t24ui*^ (n=8) and **h** *kcnj13*^*td15*^ (n=12). **i-m** Melanophore clearance in *kcnj13*^*tdxg6*^ is similar to wild type during the development of the first light stripe between 25 and 45 dpf. **n, n’** *D. rerio* wild-type and **o, o’** *kcnj13*^*tdxg6*^ patterns at J stage (11 mm). In the mutants, iridophores fail to reiterate the consecutive light stripes, which ultimately leads to fewer and broader stripes with occasional interruptions. Light and dark grey bars represent light and dark stripe areas, respectively.

In *D. rerio kcnj13* mutants fewer, wider and interrupted stripes develop, and melanophores and compact xanthophores fail to separate completely (Fig. 1c,e,f,g)^14,19,23-28^. A CRISPR/Cas9-mediated loss-of-function allele of *kcnj13* in *D. aesculapii* showed that the gene is also required for the formation of vertical bars in this species. This null allele leads to a complete loss of any pattern with uniform distribution of mixed pigment cells in the skin (Fig. 1d)^23^. Hybrids between the two species display stripes similar to the pattern in *D. rerio*. The evolutionary divergence of *kcnj13* between *D. rerio* and *D. aesculapii* was demonstrated by reciprocal hybrids between wild-type and mutant fish^23^. This genetic test is used to identify evolved genes by comparing the phenotypes of reciprocal hemizygotes; that is hybrids, which carry a null allele from either one of the parental species in an otherwise identical genetic background^29^. It depends on the ability to generate null alleles in a given species pair, which is possible in several *Danio* species since the introduction of the CRISPR/Cas9 system. Hemizygous hybrids between *D. rerio kcnj13* mutant and *D. aesculapii* wild type display a spotted phenotype indicating that *D. aesculapii* allele fails to complement the *D. rerio* null-allele, whereas the reciprocal hybrid in which the *D. aesculapii* allele was mutant displayed the striped phenotype of hybrids between the wild-type species. The different phenotypes demonstrated that the wild-type alleles from the two species are functionally no longer equivalent. Mutations in *gja4, gja5b* and *igsf11* in *D. aesculapii* revealed functions for all these genes in the formation of the bar pattern. However, all hemizygous hybrids showed patterns indistinguishable from patterns of wild-type hybrids, ruling out functional evolution of these loci. Hybrids between *D. rerio kcnj13* mutants and seven additional *Danio* species suggest that *kcnj13* evolved independently several times in the genus, as the wild-type alleles from three different species, *Danio aesculapii, Danio tinwini* and *Danio choprae*, do not complement a *D. rerio kcnj13* loss-of-function allele in hemizygous fish^23^.

In chimeras produced by blastula transplantations, we corroborate previous studies^19,25^ showing that *kcnj13* function is cell-autonomously required in melanophores but not in xanthophores for normal stripe formation. In addition, we show that the gene function is also not required in iridophores, the third pigment cell type. In vitro experiments have shown that the function of *kcnj13* is required for the depolarization of melanophore membranes upon contact with xanthophores^26^. This form of contact-dependent depolarisation might underlie the repulsive interactions between melanophores and xanthophores during the establishment of the striped pattern. To test the effects of *kcnj13* loss-of-function on the shapes of pigment cells in vivo we performed further blastula transplantations, fluorescence imaging of labelled pigment cells and cell-lineage tracing of marked clones. We find that the shapes of all three types of pigment cells are altered in the mutants, suggesting that cell-cell interactions responsible for the location-dependent acquisition of cell shapes are dependent on *kcnj13* function and defective in the mutants. Using a newly generated CRISPR/Cas9-mediated knock-in reporter line we detect *kcnj13* expression in only very few differentiated melanophores in the skin, suggesting that *kcnj13* function might be required only during a short period or in a subset of cells for a longer time during pattern formation.

The coding sequence for *kcnj13* is highly conserved within the *Danio* genus with very few non-synonymous changes between the species. However, it was not clear whether these changes between *D. rerio* and *D. aesculapii* are functionally relevant, or whether cis-regulatory evolution underlies *kcnj13* divergence^23^. We show that transgenic rescue of the *kcnj13* mutant phenotype is possible with the wild-type coding sequences of both, *D. rerio* and *D. aesculapii*, suggesting that both proteins are functionally equivalent. Strikingly, we observe a much higher expression of the *D. rerio* allele compared to the *D. aesculapii* allele in the skin of wild-type hybrids. We conclude that regulatory rather than protein changes underlie the evolution of the gene between *D. rerio* and *D. aesculapii*. The differences in the two patterns might result in part from the lower expression of *kcnj13* in *D. aesculapii* leading to variation in pigment cell distribution and shapes reminiscent of those in *D. rerio* mutants deprived of *kcnj13* activity.

## Results

### Development of the *kcnj13* phenotype in *D. rerio*

To understand the function of *kcnj13* during pattern formation, we focused on its role during stripe formation in *D. rerio*. Multiple dominant alleles of *kcnj13* have been found in several independent genetic screens^14,19,23-28^. Fish homozygous for two dominant alleles (Fig. 1e,f) and homozygotes for a recessive loss-of-function allele (Fig. 1c) develop similar but variable phenotypes with fewer, wider and interrupted stripes. To test whether this variability in our stocks is attributable to the nature of the allele (dominant or recessive) or the genetic background, we compared different allelic combinations in F2 fish with the same genetic background and found that all of them lead to indistinguishable phenotypes. This indicates that dominant and recessive alleles cause the same developmental effects in homozygous mutants (Fig. 1g,h) showing that the dominant alleles are dominant-negative in heterozygotes.

We followed the development of the mutant pattern during metamorphosis. As previously described^25^, and comparable to wild type, melanophores in the mutants are cleared from the region of the first light stripe, where compact iridophores and xanthophores develop (Fig. 1i-m). However, unlike in wild-type fish, iridophores later fail to initiate the consecutive light stripes, which leads to a phenotype of fewer and broader stripes in the mutants with occasional interruptions (Fig. 1n,o).

### Cell-autonomy of the *kcnj13* function in *D*. *rerio*

Melanophores but not xanthophores require *kcnj13* function for stripe formation as shown in chimeras created by blastula transplantations^25^. We confirmed these findings and also tested the requirement of *kcnj13* in iridophores. In these experiments the donor embryos were mutant for *kcnj13* and genetically able to provide only one of the three pigment cell types. Hosts were wild-type for *kcnj13* but lacking this pigment cell type. Thus, in three sets of transplantations, the resulting chimeras had one mutant pigment cell type placed adjacent to the respective other two wild-type cell types. In contrast to mutant xanthophores and iridophores, only mutant melanophores could not contribute to wild-type patterns in chimeras (Fig. 2a-c) leading to the conclusion that *kcnj13* is cell-autonomously required in melanophores but not in xanthophores or iridophores. By transplanting *kcnj13* mutant cells into *albino*/*slc45a2* hosts we further tested whether mutant melanophores can integrate into a normal pattern with wild-type melanophores in the chimeric animals. We observed disruptions in the striped pattern wherever mutant (pigmented) melanophores were present (Fig. 2d). Similar severe pattern defects were never observed in chimeras that had not received mutant melanophores suggesting the absence of any functional requirement in non-pigment cells (Fig. 2d). These results indicate that stripe formation requires *kcnj13* function autonomously only in melanophores or their progenitors.

**Fig. 2:**
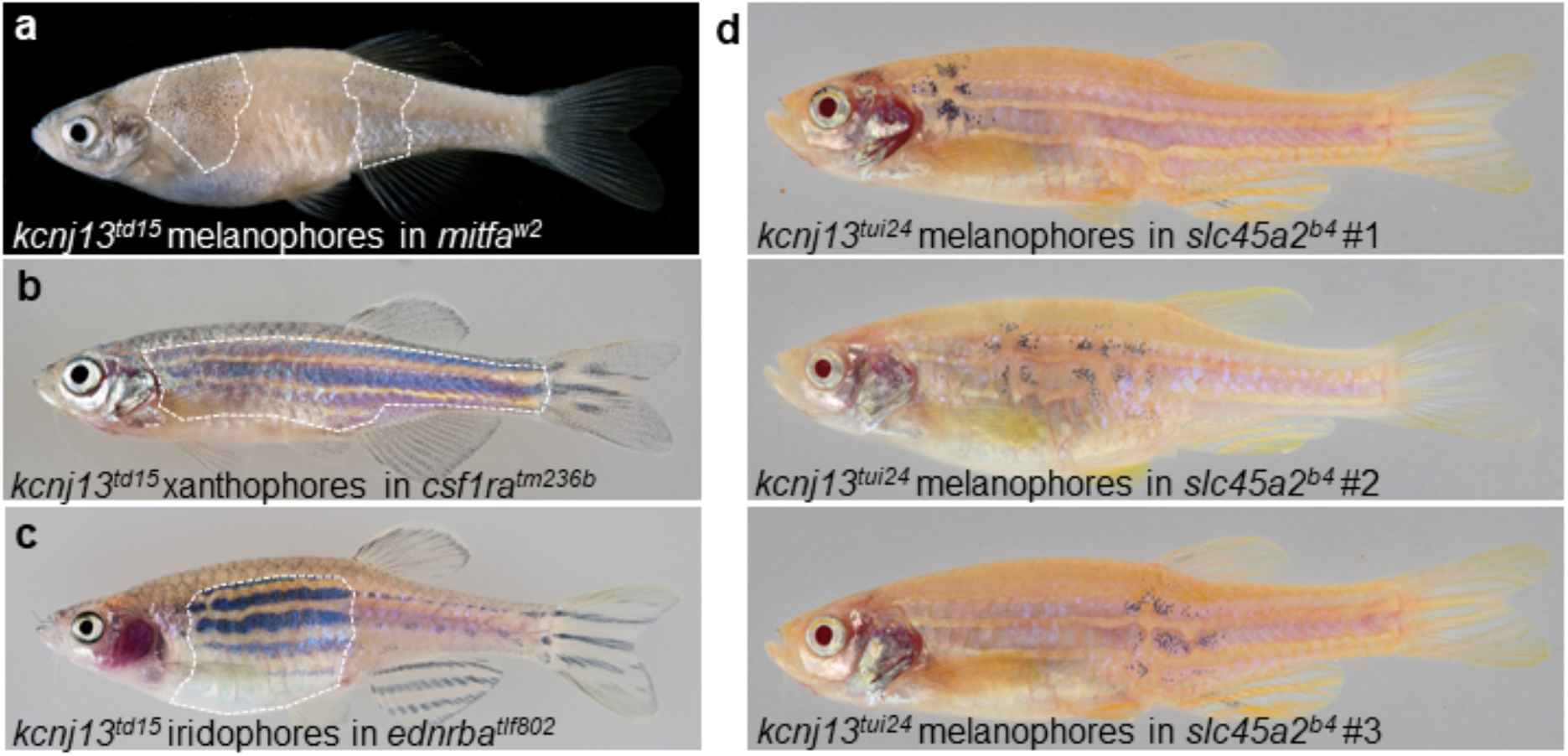
Melanophores require *kcnj13* autonomously during stripe formation. **a** Testing cell-autonomy of *kcnj13* by blastula transplantations reveals a genetic requirement in melanophores (*kcnj13*^*td15*^*;ednrba*^*tlf802*^*;csf1ra*^*tm236b*^ into *mitfa*^*w2*^), but not in **b** xanthophores (*kcnj13*^*td15*^; *kita*^*b134*^*;ednrba*^*tlf802*^ into *csf1ra*^*tm236b*^) or **c** iridophores (*kcnj13*^*td15*^*;mitfa*^*w2*^*;csf1ra*^*tm236b*^ into *ednrba*^*tlf802*^). **d** Transplantation experiments (*kcnj13*^*tui24*^ into *slc45a2*^*b4*^) provide further evidence of a cell-autonomous function of *kcnj13* in melanophores during stripe formation. Transplanted mutant melanophores (pigmented) are associated with stripe perturbations in *albino* hosts (n=3). Strong pattern deformations are never observed in chimeras without pigmented trunk melanophores (n=41).

### Endogenous *kcnj13* expression during metamorphosis in *D*. *rerio*

To investigate when *kcnj13* functions in the melanophore lineage, we used CRISPR/Cas9-mediated homology-directed repair to produce a KalTA4::Venus knock-in line (for details see methods) as a reporter for endogenous *kcnj13* expression in *D. rerio*. In early larvae we observed expression in the pronephros, hindbrain and melanophores, a pattern very similar to previously published results obtained by in situ hybridization^28^, suggesting that our reporter line faithfully recapitulates endogenous *kcnj13* expression (Fig. 3a). During later stages, at the onset of metamorphosis, expression is detected in patches of cells in the spinal cord along the entire anterior posterior axis of the fish (Fig. 3b). These positions do not overlap with the DRGs, where the neural-crest derived stem cells for the pigment cells are located^7-9^ (Fig. 3c). We conclude that *kcnj13* does not provide a function for stripe formation in these cells as our transplantation experiments indicate no functional requirement in non-pigment cells (Fig. 2d). While the signals in the kidney and spinal cord persist throughout metamorphosis, we do not find expression of the reporter in pigment cell progenitors, but in a few xanthophores and melanized melanophores in the skin during the time of pattern formation (Fig. 3d-f). These results show that *kcnj13* is expressed at detectable levels only in a small subset of melanophores at any given time during pattern formation.

**Fig. 3:**
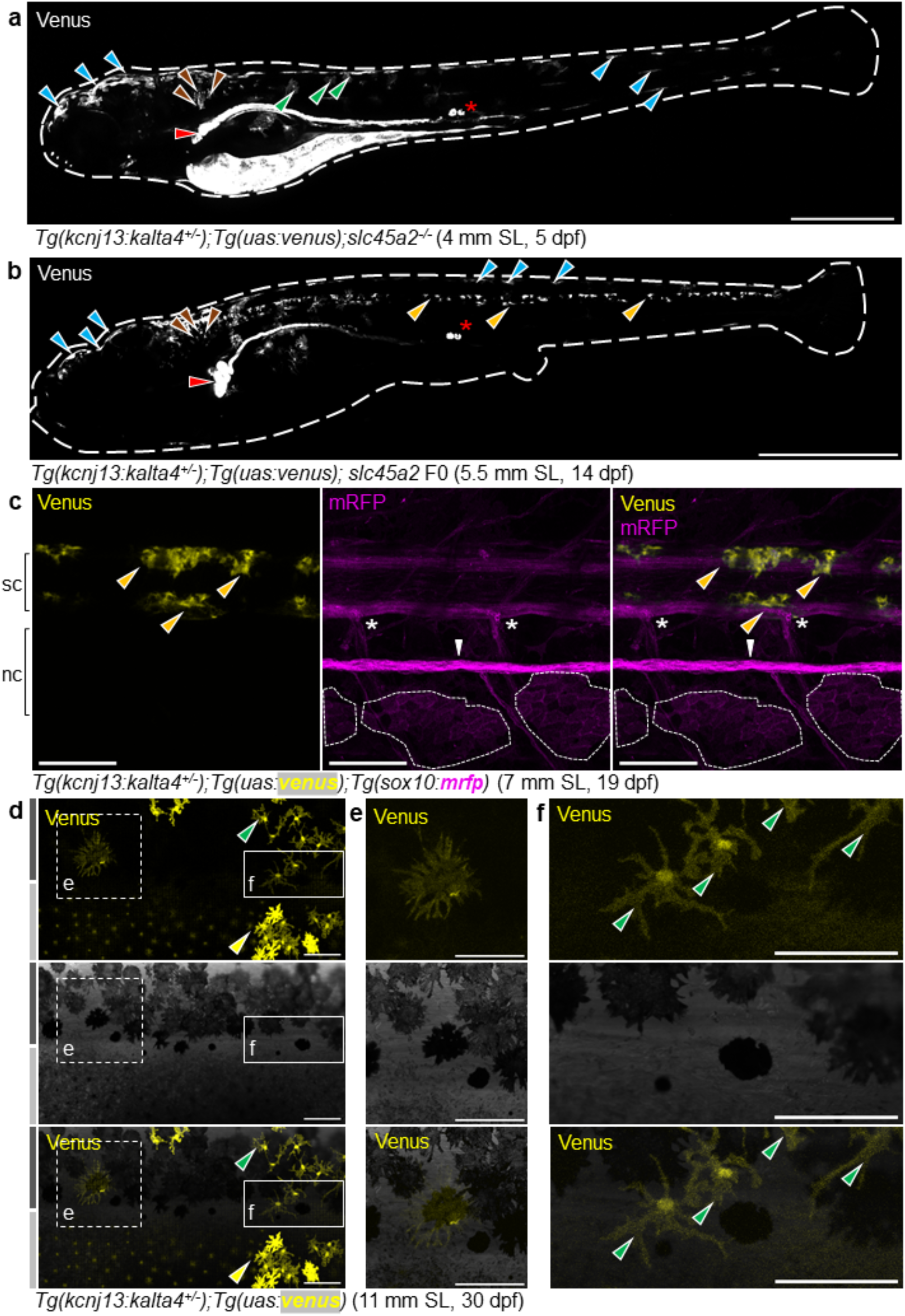
Endogenous *kcnj13* expression during *D. rerio* development. **a** Heterozygous KalTA4::Venus reporter larva showing signals in melanophores in the head and tail regions (cyan arrowheads), xanthophores (green arrowheads), hindbrain (brown arrowheads), along the entire pronephros (red arrowhead), including corpuscles of Stannius (red asterisk), and the yolk. 4 mm SL, 5 dpf, sagittal view, images of four positions along the AP axis combined into one composite; scale bar=500 µm. **b** Similar expression patterns can be observed in larva week older, with additional signals in the spinal cord (orange arrowheads). These signals persist throughout further development. 5.5 mm SL, 14 dpf, sagittal view, images of five combined into one composite; scale bar=1 mm. **c** Venus expression does not overlap with locations of the pigment cell stem cells at the DRGs (marked by white asterisks). Iridophore patches in the skin indicated with white dashed circles, lateral line nerve marked with a white arrowhead. nc: notochord, sc: spinal cord; 7 mm SL, 19 dpf; scale bar=100 µm. **d** During and after the consolidation of the stripes in wild types (see Fig. 1i-o), Venus expression can be detected in only a minority of **e** melanophores and **f** xanthophores in the skin at any given time point. Green arrowheads indicate stellate and Venus-positive xanthophores in the dark stripe, while yellow arrowheads indicate compact, pigmented and Venus-positive xanthophores in the light stripe. 11 mm SL, 30 dpf, scale bar=100 µm.

### Effects of *kcnj13* mutations on pigment cell shape in *D*. *rerio*

A key aspect of pigment pattern formation in *D. rerio* is the location-specific acquisition of different pigment cell shapes. In the dark stripes of wild-type *D. rerio*, melanophores are densely packed and compact, only cells located at the boundaries to the light stripes form long protrusions, possibly interacting directly with xanthophores and iridophores^10,30^. To investigate cell shapes in *kcnj13* mutants we observed fish carrying *Tg(kita::mcherry)*, which labels both xanthophores and melanophores. Some cells are unlabelled due to the variegation of the transgene, which allows to visualize the shapes of the tightly packed melanophores. Similar to previous findings^19^ we observed that in the dark regions in the mutants melanophores are less compact and less tightly packed compared to wild-type cells. We also find that the melanophores bordering the light stripes lack the very long protrusions present in the wild type (Fig. 4a,b). This suggests that *kcnj13* mutant melanophores do not interact with one another and with xanthophores and iridophores in the same way wild-type melanophores do.

**Fig. 4:**
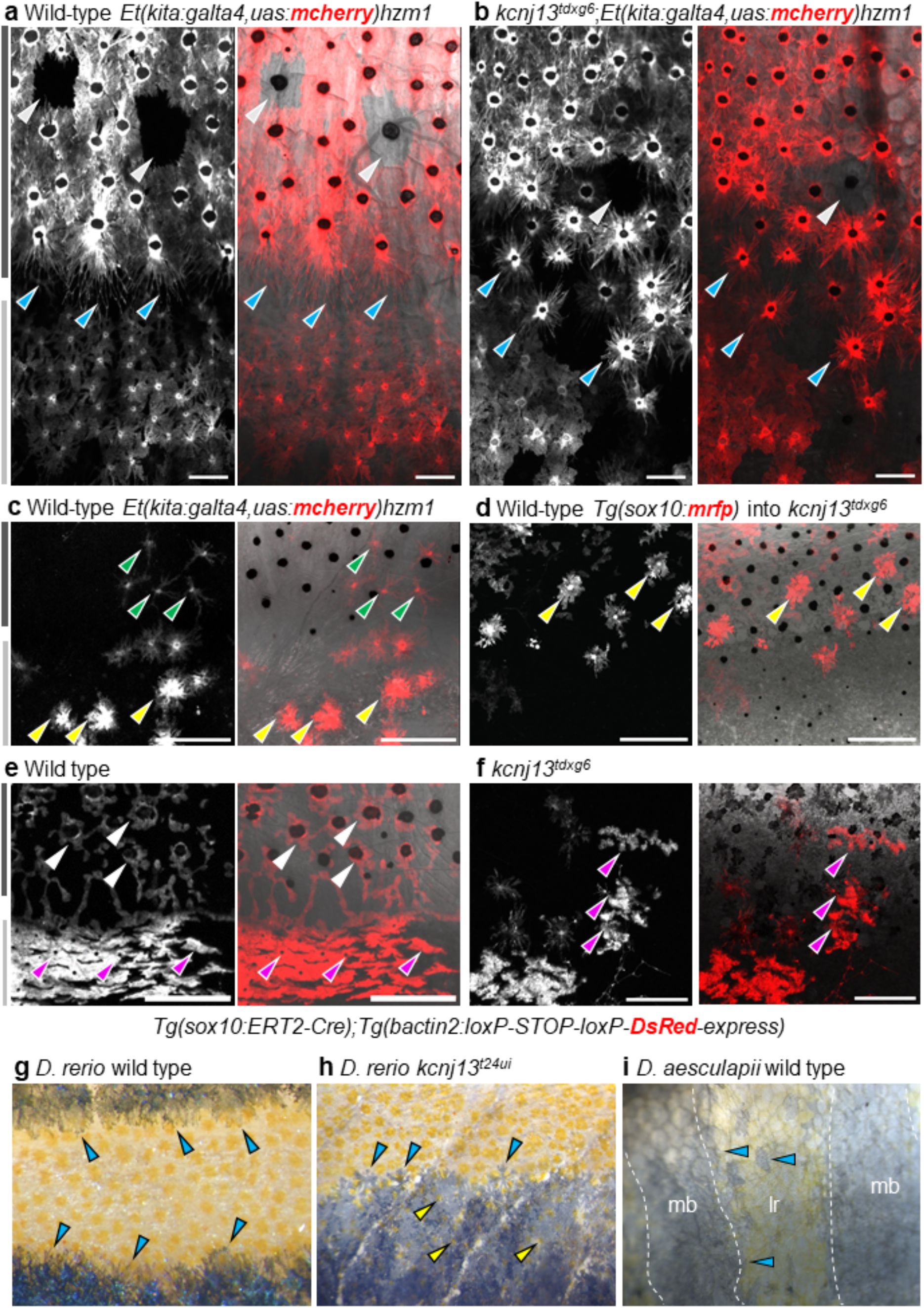
Pigment cell organization and shapes in *D. rerio* wild types and *kcnj13* mutants, and *D. aesculapii* wild types. **a** In adult wild-type *D. rerio*, melanophores in the stripe are densely packed (note variegation of the transgene in a few cells indicated with light-grey arrowheads) and cells at the boundary form long protrusions towards the light stripe (cyan arrowheads). **b** In *kcnj13*^*tdxg6*^ mutants, cells are less tightly packed in the dark stripe and short protrusions form without clear polarity (cyan arrowheads). **c** Wild-type xanthophores acquire stellate shapes in the dark stripes (green arrowheads) and compact shapes in the light stripes (yellow arrowheads). **d** Transplanted mRFP-positive wild-type xanthophores acquire inappropriate compact shapes (yellow arrowheads) in a dark stripe in *kcnj13*^*tdxg6*^ mutants (donor: *Tg(sox10:mrfp)*, host: *kcnj13*^*tdxg6*^). **e** Wild-type iridophores acquire loose shapes (white arrowheads) in the dark stripes and dense shapes (magenta arrowheads) in the light stripes. **f** Iridophores acquire ectopic compact shapes (magenta arrowheads) in the dark stripes in *kcnj13*^*tdxg6*^ mutants, visualized by tracing labelled clones. Light and dark grey bars represent light and dark stripes in *D. rerio*, respectively. **g** Wild-type *D. rerio* form long melanophore protrusions towards the light stripe regions (cyan arrowheads, see a). **h** Melanophore protrusions are not polarized in *D. rerio kcnj13* mutants (cyan arrowheads, see b) and pigmented xanthophores are visible in the dark stripe region (yellow arrowheads). **i** *D. aesculapii* wild types lack polarized melanophores (cyan arrowheads), melanophores and xanthophores mix occasionally, and the boundary between bars and light regions is of very low contrast. mb=melanophore bar region, lr=light region.

Next, we investigated the effect of *kcnj13* mutations on xanthophore behaviour during stripe formation. Upon transplanting wild-type xanthophores, labelled with *Tg(sox10:mrfp)*, into *kcnj13* mutants these cells acquire compact shapes in the dark stripe regions, where they normally appear stellate (Fig. 4c,d). Similar to findings from in vitro studies^26^, these results suggest that wild-type xanthophores are not always able to interact with mutant melanophores, which causes patterning defects in vivo.

To assess the effects of mutations in *kcnj13* on iridophores, we induced fluorescently labelled clones in the mutants using a *Tg(sox10:cre-ERt2*) line^7^ and followed labelled iridophores during metamorphosis. We found clones of dense iridophores, which are characteristic for light stripes, in the dark stripe area (Fig. 4e,f). This result suggests hat iridophores require the presence of and interaction with melanophores to acquire the loose form; and that this interaction depends on *kcnj13* function. Thus, iridophores might not be able to recognise mutant melanophores and therefore develop ectopically in the dense form in the dark stripe regions. We conclude that *kcnj13* function, required in melanophores, is important for homotypic and heterotypic pigment cell interactions, which control the location-dependent cell shape acquisition of all three pigment cell types during pattern formation. These cumulative effects might inhibit the reiteration of dark and light stripes in the mutant fish.

### Evolution of pigment cell shapes between *D*. *rerio* and *D*. *aesculapii*

Melanophores in *D. rerio* produce pronounced polarized protrusions towards compact xanthophores and both cell types are strictly separated between the light and dark stripes. The polarity of the protrusions is lost in *kcnj13* mutants, where both cell types also mix occasionally (Fig. 4a,b,g,h). This mutant phenotype is similar to the situation in wild-type *D. aesculapii*, where we found a mixing of cells and no pronounced polarity of melanophores towards xanthophores (Fig. 4i). The contrast of the bar pattern is therefore reduced; there is no contrast in *D. aesculapii kcnj13* mutants, where all pigment cells mix and no bars are formed^23^. Our observations suggest that the divergence of the pigment patterns between *D. rerio* and *D. aesculapii* could partially be due to evolutionary changes in the interactions between all three pigment cell types, which influence the cell shapes.

### Molecular basis of *kcnj13* evolution between species

To investigate the channel structure of Kcnj13 (Kir7.1), we expressed the *D. rerio* protein fused to mCherry using a Multibac-derived baculovirus/insect cell expression system^31,32^, purified the recombinant protein by affinity and size-exclusion chromatography, and measured the molecular mass with mass photometry^33^ (Supplementary Fig. 1). The results suggest that Kcnj13 exists as a homo-tetramer, which can explain the dominant-negative effects observed in alleles carrying point mutations affecting the selectivity filter or the second transmembrane helix (Fig. 1a,b) as caused by mutant proteins negatively interfering with wild-type copies in the complex in heterozygous fish^14,19,23,24,27,28^. We constructed homology-based and AlphaFold-multimer models of the homo-tetrameric Kcnj13 channel (Supplementary Files). These models agree with published structures of similar potassium channels. The protein sequences of *D. rerio* and *D. aesculapii* differ only by two amino acid residues (Q23L and D180G in magenta) in the cytoplasmic domain^23^; structure modelling of the two alleles is insensitive to these differences.

Reciprocal hemizygosity tests showed that the divergence of *kcnj13* must reside within the locus, either in the protein-coding region or in cis-regulatory elements, but cannot be due to trans-acting factors^23^. To test whether the amino acid changes identified between the two species contribute to the evolution of *kcnj13*, we used Tol2 transgenesis to express the coding regions from *D. rerio* or *D. aesculapii* under the control of the melanophore-specific *mitfa* promoter in *kcnj13* null-mutant *D. rerio* (Fig. 5b,d). In both cases the transgenes were able to restore the striped pattern in the trunk of the fish, indicating that the protein from *D. aesculapii* can function in a similar manner to the *D. rerio* protein (Fig. 5e). We observed some differences in the rescue capabilities of the transgenes among the lines we established, possibly due to copy number variations and expression differences of the randomly inserted transgenes. The striped pattern of the caudal fin was never restored in the transgenic lines, most likely due to the inactivity of the promoter at the appropriate time points in this tissue, corroborating the finding of fundamental mechanistic differences in pigment pattern formation between the trunk and fin (Frohnhöfer et al. 2013). Our results suggest that the coding regions from both species function similarly and that the protein-coding changes are irrelevant for *kcnj13* divergence.

**Fig. 5:**
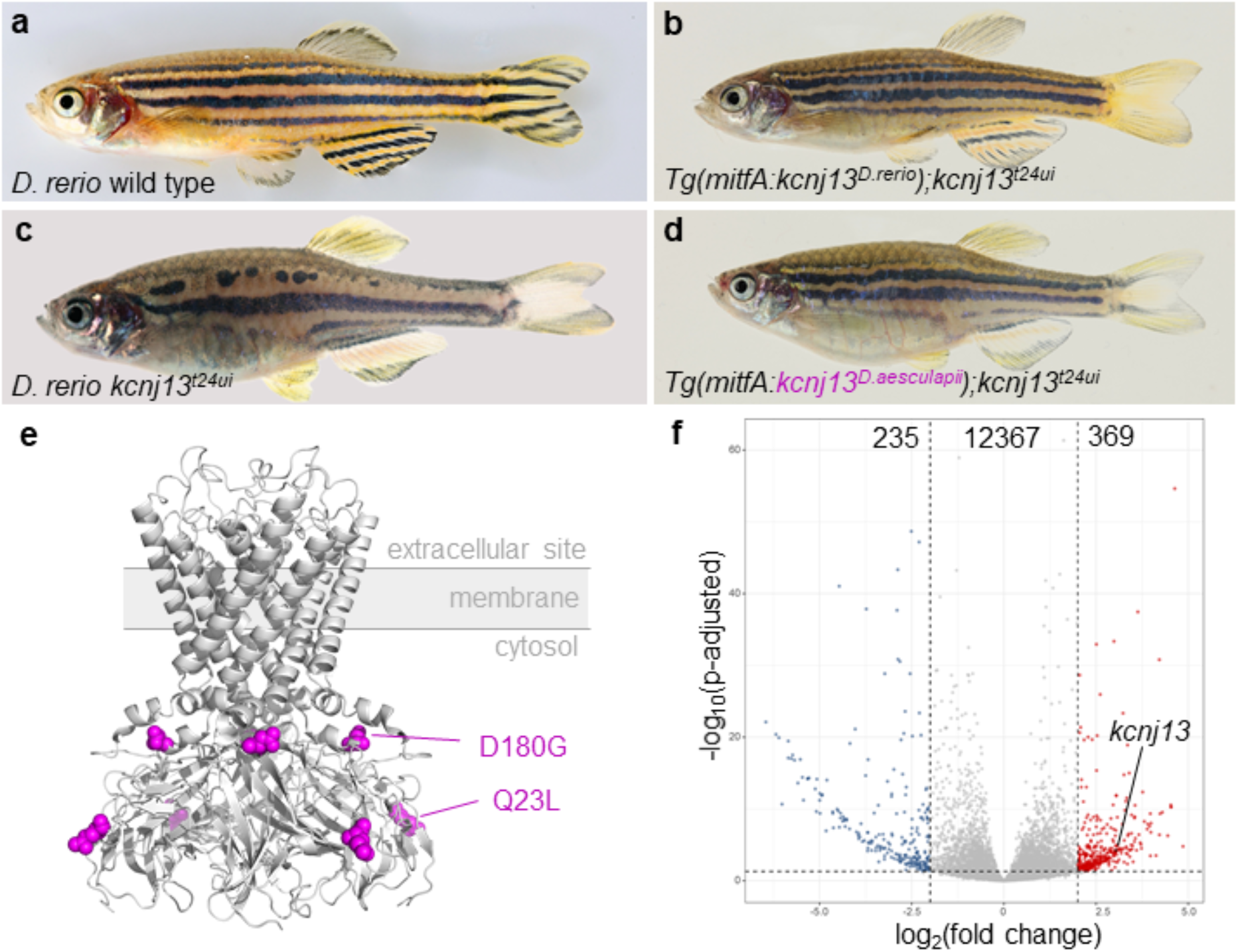
Molecular basis of *kcnj13* evolution between *D. rerio* and *D. aesculapii*. **a** *D. rerio* wild type. **b** *D. rerio kcnj13*^*t24ui*^, in which either **c** the *D. rerio* allele of *kcnj13* (*Tg(mitfA:kcnj13*^*D*.*rerio*^*);kcnj13*^*tui24*^) or **d** the *D. aesculapii kcnj13* allele (*Tg(mitfA:kcnj13*^*D*.*aesculapii*^*);kcnj13*^*tui24*^) was expressed under the control of the *mitfa* promoter from *D. rerio*. In both cases, stripes were restored in the trunk of the fish. R224K was found to be polymorphic in *D. aesculapii* (Podobnik et al. 2020). **e** SWISS-MODEL derived homology model of the Kcnj13 tetramer (Q23L and D180G diverged between species in magenta). **f** Allele-specific expression analysis in interspecific hybrids shows higher *kcnj13* expression of the *D. rerio* allele in the skin (n=12; *p-adjust* < 0.0001), confirming cis-regulatory evolution. Overall, we found no differences in expression levels in 12,367 genes. 369 and 235 genes were significantly higher expressed from the *D. rerio* (red) or *D. aesculapii* allele (blue), respectively.

Therefore cis-regulatory changes likely underlie *kcnj13* evolution and patterning differences between the two species. To test this prediction, we produced hybrids between the two species and performed allele-specific expression analysis in the skin and posterior trunk of adult fish. We found significantly higher levels of the *D. rerio* allele compared to the *D. aesculapii* allele (Fig. 5f, Supplementary Fig. 2), indicating species-specific regulation of the locus and thereby confirming cis-regulatory evolution. Quantitative differences in expression levels might cause differences in pigment cell interactions and shapes observed between *D. rerio* and *D. aesculapii*. Based on the repeated and independent evolution of the ancestral *kcnj13* function in the *Danio* genus^23^ we speculate that similar cis-regulatory changes might also have occurred in *D. tinwini* and *D. choprae*. Our results highlight the *Danio* genus as an excellent model system to study the molecular, genetic and cellular basis of pigment pattern diversification in vertebrates.

## Discussion

Teleost fish produce some of the most intricate pigmentation patterns in nature. However, only in a few species the pattern forming mechanisms are studied in detail. *D. rerio*, an excellent vertebrate model organism widely used in research, shows a conspicuous pattern of horizontal stripes on the flank and in the anal and tail fins. This pattern is produced by three types of pigment cells interacting in complex ways to self-organize into dark and light stripes. During pattern formation the horizontal myoseptum serves as an anatomical pre-pattern for the orientation of the stripes. The stripes in the anal and tail fins are contiguous with the stripes in the body. However, the fin pattern is formed by a different, possibly somewhat simpler mechanism that involves only two cell types, melanophores and xanthophores. Cellular interactions mediated by direct cell-cell contacts depending on gap junctions and adhesion molecules are essential for stripe formation as demonstrated by the spotted phenotypes of *gja4, gja5b, igsf11* and *jam3b* mutants^14,15,17,18^. In addition, mutations in *kcnj13* lead to defects in the pattern with fewer, wider and interrupted stripes and occasional mixing of compact xanthophores with melanophores^14,19,23,24,26-28^. Kcnj13 regulates the membrane potential of melanophores^26^, which might be important for the repulsion between xanthophores and melanophores. By interspecies complementation tests in *Danio* hybrids it was previously shown that of these four genes only the function of *kcnj13* diverged within the *Danio* genus, probably several times independently^23^.

To better understand the role of *kcnj13* in pattern formation and diversification, we examined its function in *D. rerio* in more detail. All *kcnj13* alleles isolated in genetic screens are dominant with a relatively weak heterozygous and considerably stronger homozygous phenotype. We previously produced a loss-of-function allele, which is completely recessive^23^. The phenotypes of homozygous fish for a dominant or the recessive allele in the same genetic background are indistinguishable (Fig. 1g,h). This demonstrates that the dominant alleles are in fact dominant-negatives and not neomorphs. The variability we observe in our mutant strains is dependent on the genetic background.

Phenotypic analysis of chimeras obtained by blastula transplantations had already demonstrated the autonomous requirement of *kcnj13* function in melanophores but not in xanthophores^25^. We repeated these transplantation experiments including the third pigment cell type, iridophores. Our results show that *kcnj13* function is required only in melanophores for stripe formation in *D. rerio*, but not in any other cell type (Fig. 2a-c). In addition, we find that mutant melanophores lead to strong patterning defects when transplanted into wild-type fish (Fig. 2d). This shows that the mutant cells are not guided by their wild-type neighbours but influence the patterning process cell-autonomously, possibly failing to instruct neighbouring xanthophores and iridophores.

Our results support the prior observation that a *kcnj13* transgene expressed under the control of the *mitfa* promoter, which is known to be active in melanophores and their stem cells^9^, can rescue the mutant phenotype in the trunk^26^. As these experiments were conducted in the presence of a dominant-negative *kcnj13* allele, which impedes the wild-type channel function, a complete rescue could not be expected. In our transgenic rescue experiments, using the recessive mutant, expression of *kcnj13* using the *mitfa* promoter restores the stripes on the flank of the fish to a pattern very similar to the one observed in wild types (Fig. 5b), which further supports the notion that *kcnj13* is required in melanophores. The striped pattern in the anal and tail fins is not restored by the transgenes suggesting that expression under the melanophore-specific *mitfa* promoter does not recapitulate all aspects of the endogenous expression pattern of *kcnj13*, and mechanisms that form stripes in the fins are fundamentally different from those that form stripes in the trunk^10^.

To visualize the expression pattern of *kcnj13* in *D. rerio* we made a reporter line by homology directed knock-in of an optimized GAL4 coding sequence (KalTA4) into the endogenous locus. In combination with a UAS:Venus transgene this reporter line shows expression in early larvae in the pronephros and melanophores (Fig. 3a,b), very similar to published data from in situ hybridizations^28^, indicating that our line faithfully recapitulates *kcnj13* expression. Later, during metamorphosis when the pigment pattern is formed and also in adult fish, we detected expression in neurons of the spinal cord (Fig. 4c). During these stages in situ hybridizations are difficult in *D. rerio* and we rely on the reporter to indicate expression of the gene. As our transplantation experiments clearly show a cell-autonomous requirement of *kcnj13* in melanophores or their precursors (Fig. 2d) we can rule out a function of the gene for pattern formation in these neuronal cells. We also found expression of the reporter line during later stages in few xanthophores and, unexpectedly, only in a small subset of melanophores (Fig. 4d). Expression of the reporter in xanthophores might reflect earlier activation in a common precursor for melanophores and xanthophores and the long persistence of the proteins (KalTA4 and Venus). Alternatively, *kcnj13* could genuinely be expressed in xanthophores but without any obvious function in stripe formation. Our observation that we cannot detect *kcnj13* expression in all melanophores at any given time point suggests that it is either required only very transiently or that only a few cells depend on *kcnj13* function and then influence the behaviours of all the pigment cells. Alternatively, our reporter might not be sensitive enough to allow the detection of very low expression levels, which could nevertheless be relevant for pattern formation. A different possibility is that the channel protein might be very stable and present in the cell membrane for prolonged times even after transcription has ceased and also the reporter is no longer detectable. In any case, our data is consistent with published data from single-cell RNA sequencing^34^, which also show expression of *kcnj13* to be low and limited to a very minor fraction of pigment cell progenitors as well as differentiated melanophores and xanthophores.

We conclude that *kcnj13* is only required in melanophores during pattern development. Mutant melanophores are less compact and less tightly packed affecting the tiling within the dark stripe. Mutant melanophores at the stripe boundaries also do not form polarized protrusions towards the light stripes (Fig. 4a,b). The significance of these protrusions is unclear, they could be used for direct repulsive interactions with xanthophores or iridophores to delineate the boundary between light and dark stripe^10,30^. In *kcnj13* mutants homotypic and heterotypic interactions, among melanophores and between melanophores and the other two pigment cell types, are affected, as seen, for example, by the mixing of the cells. We find that the shapes of both cell types are affected in *kcnj13* mutants, with dense iridophores and compact xanthophores, which are limited to the light stripes in wild type, also appearing in dark stripe regions. Therefore, we conclude that melanophores play a critical *kcnj13*-dependent role in directing dark stripe-specific cell shape transitions in both, iridophores and xanthophores. In the absence of Kcnj13 all three types of pigment cells may lose their dark stripe-specific shapes, which might indicate that the default shapes for xanthophores and iridophores are the ones these cells acquire in the light stripe region.

The same types of pigment cells that are found in *D. rerio* form a range of very different patterns in closely related *Danio* species. The specification and differentiation of pigment cells are similar in *D. rerio* and *D. aesculapii*. They both require Mitfa- and Kit signalling in melanophores and Csf1 and Ltk signalling in xanthophores and iridophores, respectively^23,35^. Mutants indicate that iridophores do not emerge along the horizontal myoseptum, are lower in number and dispensable for bar formation in *D. aesculapii* whereas they guide stripe formation in *D. rerio*^23^.

Whether genes required for iridophore development have evolved between these two species is not known. However, for another species, *D. nigrofasciatus*, it was shown that reduced iridophore proliferation contributes to a reduction in stripe number and integrity^22^. In addition, species-specific differences in the developmental timing of pigment cell proliferation and differentiation can lead to patterning differences as observed for xanthophores, which differentiate precociously in *D. albolineatus* resulting in a loss of the striped pattern^21^. We find that melanophores in *D. aesculapii* do not form long protrusions towards the light regions (Fig. 4g-i), which is similar to *kcnj13* mutants in *D. rerio* (Fig. 4a,b). In *D. rerio* these protrusions might partly regulate melanophore survival^30^ and the overall stability of the boundary between dark and light stripes. Similar to the *D. rerio* mutant, the lack of such protrusions in *D. aesculapii* might indicate a less robust mechanism for the consolidation of the boundary between dark bars and light regions (Fig. 4i), where melanophores and xanthophores frequently mix.

When tested in *D. aesculapii* the four genes (*kcnj13, gja4, gja5b* and *igsf11*), known to function in cell-cell interactions during stripe formation in *D. rerio*, were found to be also required to form the bar pattern^23^. Whereas residual patterns of spots or wider and interrupted stripes still form in *D. rerio* mutants, the bar pattern is completely lost in *D. aesculapii* mutants and all pigment cells intermingle and distribute evenly in the skin, a phenotype only seen in double mutants in *D. rerio*. This indicates that cellular interactions in both species occur but are more complex in *D. rerio*, which could lead to a higher robustness of the patterning mechanism in this species. Reciprocal hemizygosity tests for all four genes lead to the conclusion that there is functional conservation in three cases, *gja4, gja5b* and *igsf11*, while only *kcnj13* diverged between the two species^23^. Thus, the formation of the very different patterns of horizontal stripes and vertical bars involves the same players. Three of these, Kcnj13 and the two gap junction proteins, might be involved in an electric coupling of pigment cells, which could allow coordinated tissue-scale patterning^36^. Evolution in *kcnj13* between the two species might influence the conditions for these interactions, with the consequence of evolutionary change in patterning.

In our rescue experiments the coding sequences from both species, *D. rerio* and *D. aesculapii*, were equally able to restore stripe formation in *D. rerio kcnj13* mutants indicating functional equivalency. However, the use of a non-native promoter and possible position effects due to random integration of the transgenes might obscure subtle functional differences between the two proteins. This question could be addressed in the future by precise exchanges in the coding sequence of the endogenous locus in *D. rerio*. However, we found allele-specific differences of *kcnj13* expression in hybrids with much higher levels of expression from the *D. rerio* allele (Fig. 5f) clearly indicating regulatory differences between the loci from the two species. Therefore, the functional divergence of *kcnj13* between *D. rerio* and *D. aesculapii* is most likely caused by evolution of cis-regulatory elements affecting the levels of expression of the gene. Cis-regulatory evolution has been implicated in other cases of pattern diversification of *Danio* fish. In *D. albolineatus* the increased expression of Csf1 causes early differentiation of xanthophores leading to a loss of the striped pattern and the mixing of pigment cells^21^. In *D. nigrofasciatus* iridophore development is reduced due to cis-regulatory changes in the Edn3 gene leading to an attenuated pattern with fewer melanophores and stripes, similar to hypomorphic *D. rerio* mutants^22^. In the rare case of *D. kyathit* and *D. quagga* hybrids between the two species are fertile, which allows for quantitative trait locus (QTL) mapping. QTL analysis for differences between the spotted *D. kyathit* and the striped *D. quagga* led to the identification of a complex genetic basis for the pattern differences with multiple candidate loci, probably involving changes in a number of regulatory regions^37^. In the more distantly related cichlids bars and stripes evolved repeatedly in species endemic to the Great African Lakes. Here, QTL mapping identified regulatory changes in the gene *agouti-related peptide 2* (*agrp2*) that underly these patterning differences^38^.

In three-spine sticklebacks genome-wide association studies identified loci underlying repeated ecological adaptations in independent pairs of fresh- and saltwater populations^39^. These adaptive loci are predominantly affected by cis-regulatory changes leading to differences in gene expression in the gills^40^. In contrast, trans-acting factors independently evolved to affect gene expression in the pharyngeal tooth plate in sticklebacks^41^. It was speculated that the genetic architecture of teeth formation is less complex than the adaptations to salt handling; evolution of trans-acting factors might therefore be less pleiotropic in dental tissue compared to multifunctional gills.

Dominant mutations in *kcnj13* in *D. rerio* cause pigment pattern defects but also late-onset retinal degeneration^42,43^, similar to mutations in the human ortholog that are known to cause two rare retinal diseases^44,45^. Mutations in mice lead to lethal defects in tracheal development^46^. Due to this observed pleiotropy protein evolution might be highly constrained, favouring regulatory evolution. In general pigment patterns seem to evolve often by regulatory mutations, whereas pigmentation frequently diverges by protein changes^47^. However, constraints on regulatory evolution also exist; ectopic expression of *kcnj13* in the dermomyotome leads to a long-finned phenotype^28^. Cis-regulatory evolution in *kcnj13* specifically affecting expression in the skin is presumably non-pleiotropic and might therefore be more permissive for evolutionary change influencing pigment cell behaviour.

A basic colour-forming unit in cold-blooded vertebrates, fish, amphibians and reptiles, consists of xanthophores in the top layer, iridophores in the middle layer and melanophores in the bottom layer. Melanophores appear black in the absence of shiny iridophores and yellow-orange xanthophores on top, as in *D. rerio shady*/*ltk* or *pfeffer*/*csf1ra* mutants. Modifications of this basic arrangement of pigment cells can yield diverse colourations. By varying the mechanisms that regulate pigment cell shape and layering, differences in colour, brightness and contrast can be achieved. In this regard our study points towards *kcnj13* as a key node for evolutionary tinkering that underlies colour pattern diversification in teleosts. *D. rerio kcnj13* mutants develop light and dark stripe regions low in contrast due to pigment cells that lack location-specific shapes and colouration. Regulation of colouration by cell shape transition may point to an important mechanism employed across evolution, where layer-specific and location-specific arrangement of diverse pigment cell types leads to species-specific colouration.

## Figures

**Supplementary Fig. 1:**
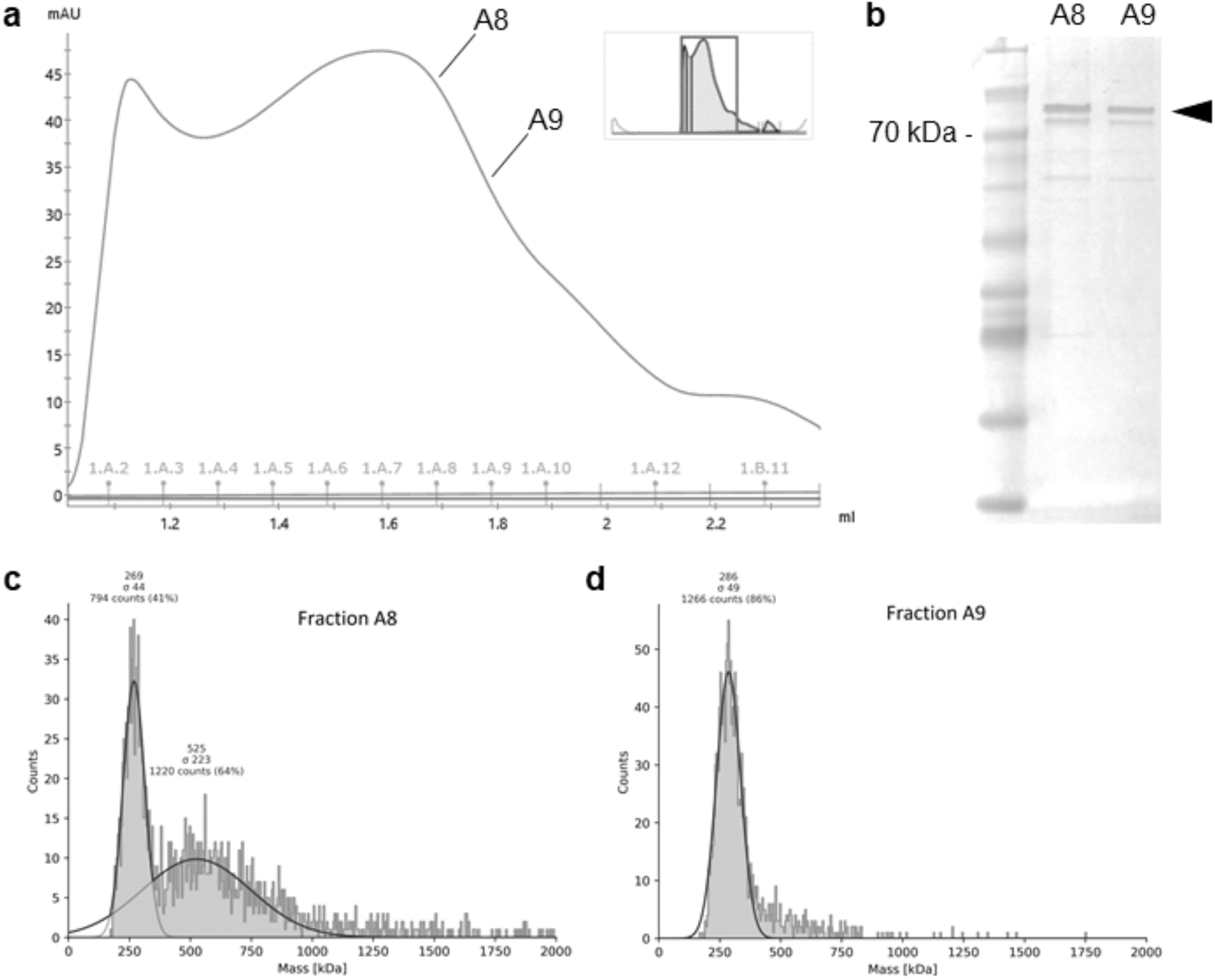
Protein purification and analysis. **a** Size-exclusion chromatogram, fractions A8 and A9 are indicated. **b** Coomassie staining shows bands corresponding to the expected size of about 70 kDa, with double bands presumably due to glycosylation. **c** and **d** show mass-photometry peaks from fractions A8 and A9, corresponding to a molecular mass of about 280 kDa, as expected for a tetrameric complex. In c higher oligomeric states might be present.

**Supplementary Fig. 2:**
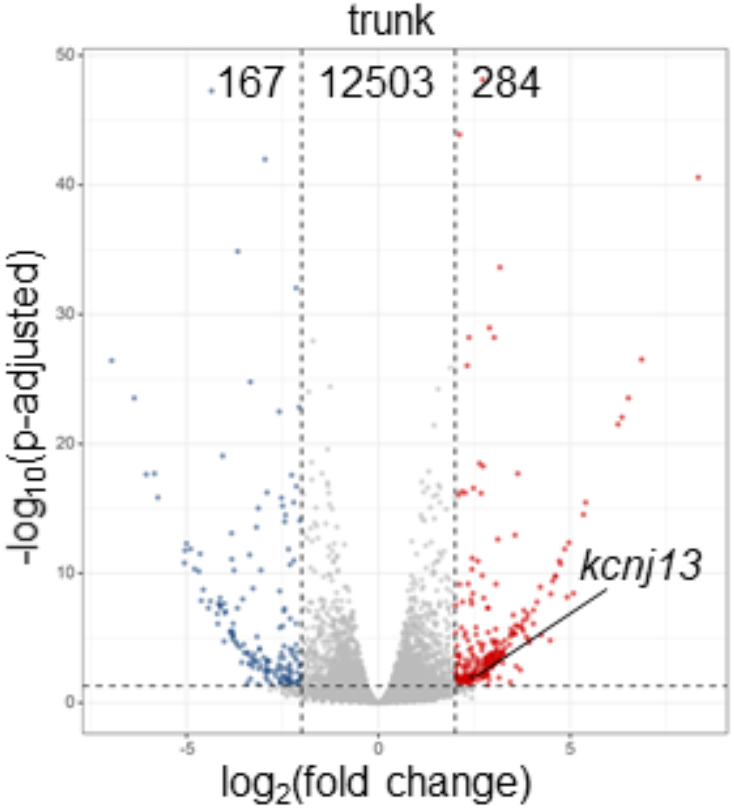
Allele-specific expression analysis in hybrids between *D. rerio* and *D. aesculapii*. In the trunk, 284 and 167 genes were significantly higher expressed from either the *D. rerio* (red) or *D. aesculapii* allele (blue). For most transcripts (12,503) we observed no differences in expression levels. We found significantly higher expression of *kcnj13* from the *D. rerio* allele (*p-adjust* < 0.05).

## Material and Methods

No statistical methods were used to predetermine sample size. The experiments were not randomized. The investigators were not blinded to allocation during experiments and outcome assessment.

### Fish husbandry

*D. rerio* and *D. aesculapii* were maintained as described in Brand & Nüsslein-Volhard^48^. If not newly generated (Supplementary Table 1), the following lines were used for experiments: *D. rerio* wild-type Tuebingen (TU), *kcnj13*^*t24ui*23^, *kcnj13*^*td15*19^, *kcnj13*^*tdxg6*14^, *nacre/mitfa*^*w2*49^, *pfeffer/csf1ra*^*tm236b*50,51^, *rose/ednrba*^*tlf802*52^, *albino/slc45a2*^*b4*53^,*sparse/kita*^*b134*54^, *Tg(sox10:mrfp)*^7^, *Et(kita:galta4,uas:mcherry)hzm1*^55^, *Tg(sox10:ERT2-Cre);Tg(bactin2:loxP-STOP-loxP-DsRed-express)*^56,57^ and *D. aesculapii kcnj13*^*tmp11*23^. Interspecific hybrids between *D. rerio* and *D. aesculapii* were obtained by in vitro fertilizations^20^. All species were staged according to the normal table of *D. rerio* development^58^. All animal experiments were performed in accordance with the rules of the State of Baden-Württemberg, Germany, and approved by the Regierungspräsidium Tübingen.

**Supplementary Table 1:**
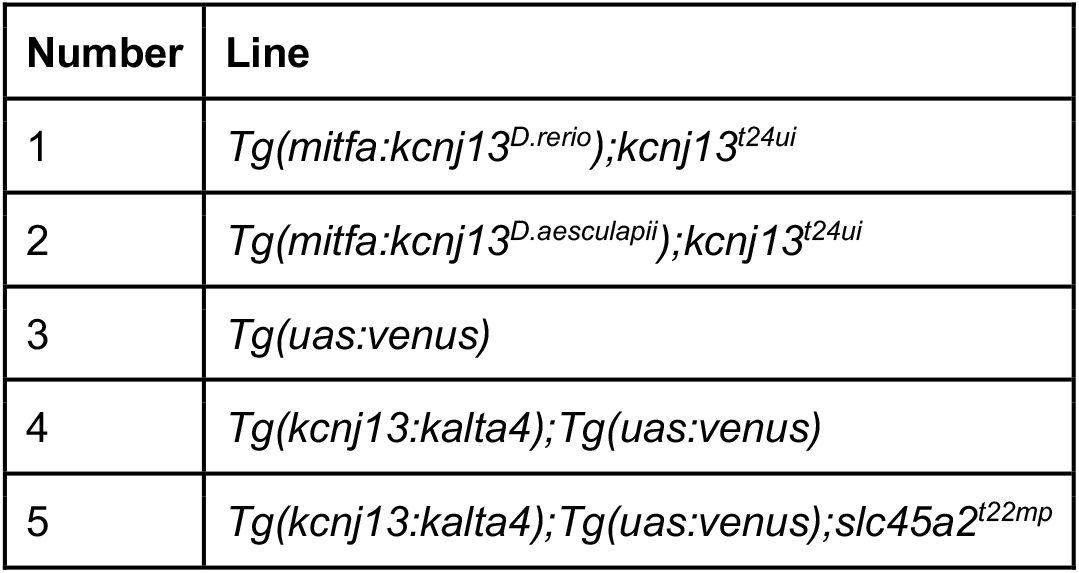
New transgenic lines used in this study.

### Tol2-mediated transgenesis

To generate the transgenic rescue lines plasmids with the *mitfa* promoter sequence from *D. rerio*^*49*^, the coding sequences of *kcnj13* from *D. rerio* or *D. aesculapii*, and the coding sequence of sfGFP was constructed. The construct was subcloned into the Tol2 vector *pGEM-T pminiTol2* carrying SV40 elements, a green heart marker *cmlc2*:*venus* and Tol2 restriction sites^59,60^. The resulting plasmids were designated as *pTol2gh-mitfa-kcnj13*^*D*.*rerio*^*-sfGFP* (GenBank accession number: OP326275) and *pTol2gh-mitfa-kcnj13*^*D*.*aesculapii*^*-sfGFP* (GenBank accession number: OP326276). Tol2 transgenesis was performed as previously described^60^; briefly, a solution (12.5 ng/µL Tol2 mRNA, 50 ng/µL plasmid DNA, and 5 % Phenol Red) was injected into fertilised eggs of *D. rerio kcnj13*^*t24ui*^ at the one-cell-stage. 100 F0 embryos were selected for marker gene expression at around 2 dpf and raised to adulthood. Mature F0 founder fish were outcrossed to *D. rerio kcnj13*^*t24ui*^ and F1 larvae positive for marker gene expression were selected to obtain stable transgenic lines. In both cases, lines were identified in which the mutant phenotype was partially rescued. These lines were designated as *Tg(mitfa:kcnj13*^*D*.*rerio*^*);kcnj13*^*t24ui*^ and *Tg(mitfa:kcnj13*^*D*.*aesculapii*^*);kcnj13*^*t24ui*^, outcrossed to *D. rerio kcnj13*^*t24ui*^, and selected for marker gene expression in embryos and intact stripe patterns in adults for at least three generations (Supplementary Table 1).

To generate a *D. rerio* UAS:Venus line a plasmid with the coding sequence for the Venus-variant of YFP under the control of the yeast transcription factor GAL4 (6 UAS-sites) was constructed (pminiTol2_UAS:Venus, GenBank accession: OP243708); mRNA for the Tol2 transposase was transcribed in vitro from the plasmid pCS2FA-transposase^61^ using the mMessageMachine and Poly-A tailing Kits (Invitrogen). TU embryos at the one-cell stage were injected with approximately 2-4 nL of injection mix containing 250 ng/µL of in vitro transcribed mRNA and 25 ng/µL of plasmid DNA in PBS with Phenol Red as a tracer dye. The adult F0 fish were crossed to TU and the F1 larvae were screened for expression of the mCherry marker in the heart. From the positive F1 fish a stable line was established by another outcross to TU followed by sibling matings of the F2 fish (Supplementary Table 1).

### CRISPR/Cas9-mediated knock-out and knock-in

For gene knock-outs the CRISPR/Cas9 system was applied either as described in Irion et al.^62^ or according to the guidelines for embryo microinjection of Integrated DNA Technologies (IDT). Briefly, oligonucleotides were cloned into pDR274 to generate the sgRNA vector. sgRNAs were transcribed from the linearised vector using the MEGAscript T7 Transcription Kit (Invitrogen). Alternatively, target-specific crRNAs and universal tracrRNAs were purchased from IDT. Cas9 was expressed as a fusion protein with mCherry in *E. coli* (BL21(DE)3pLysS) from the plasmid pOPT-Kan_Cas9-mCherry (GenBank accession: OP243709) and purified via double affinity chromatography (His-Tag and Twin-StrepTag) using standard procedures. Before use, the purified protein was dialyzed into PBS containing additionally 300 mM NaCl and 150 mM KCl, aliquoted and stored at −70°C. sgRNAs or crRNA:tracrRNA duplexes were injected as ribonucleoprotein complexes with Cas9 proteins into one-cell stage embryos. The efficiency of indel generation was tested on eight larvae at 1 dpf by PCR using specific primer pairs and by sequence analysis as described previously^63^. The remaining larvae were raised to adulthood. Mature F0 fish carrying indels were outcrossed. Loss-of-function alleles in heterozygous F1 fish were selected to establish homozygous or trans-heterozygous mutant lines (Supplementary Table 1).

To generate a reporter line for the expression of *kcnj13* the CRISPR/Cas9-system was used. For the sgRNA template two oligonucleotides (5’-TAGGCCGTCTTTGCTGACCAGG-3’ and 5’-AAACCCTGGTCAGCAAAGACGG-3’) were annealed and cloned into pDR274; the RNA was transcribed in vitro with the MegaScript Kit from Invitrogen. A donor plasmid was constructed containing the KalTA4 variant^55^ of the GAL4 coding sequence flanked by homology arms and CRISPR target sites (GenBank accession: OP243710). This plasmid (25 ng/µL) was co-injected with Cas9 protein (500 ng/µL) and sgRNA (35 ng/µL) into one-cell stage embryos from the UAS:Venus line. The resulting F0 fish were backcrossed to UAS:Venus and the F1 larvae were screened for expression of Venus. One founder fish was identified with offspring showing a very strong early signal in the yolk and later also in the pronephros and melanophores, consistent with published expression data (Supplementary Table 1). To achieve good imaging conditions in this line we generated an *albino* loss-of-function allele, *slc45a2*^*t22mp*^, as previously described^62^ (Supplementary Table 1).

### Blastula transplantations

Chimeric animals in Fig. 2a-d and Fig. 4d were generated by transplantations of cells during blastula stage as described in^64^.

### Cre induction and clonal analyses

Cre induction was carried out as described in^7^. Labelled clones in Fig. 4e,f were from fish followed over pattern development.

### Image acquisition and processing

Anesthesia of postembryonic and adult fish was performed as described previously^7^. Bright-field images of adult fish in Fig. 1a-h and Fig. 2a-d were obtained using a Canon 5D Mk II camera. To visualize melanophore protrusions via dispersion of melanosomes using bright-field imaging (Fig. 4g-i), fish were kept in the dark with a final concentration of 100 µM yohimbine (CAS: 65-19-0, Sigma-Aldrich) for 30 minutes before imaging as described in^30^. Fish with different pigment patterns vary considerably in contrast, thus requiring different settings for aperture and exposure time, which can result in slightly different colour representations in the pictures.

Fluorescence images of postembryonic and adult fish were acquired on a Zeiss LSM 780 NLO confocal (BioOptics Facility, Max Planck Institute for Biology Tübingen) and a Leica M205 FA stereo-microscope. Repeated imaging of pigment cell clones in metamorphic *D. rerio* was performed as described in^7^. Maximum intensity projections of confocal scans were uniformly adjusted for brightness and contrast. Images were processed using Adobe Photoshop, Adobe Illustrator CS6 and Fiji^65^.

### Protein expression and purification

We expressed Kcnj13-mCherry with N-terminal His-tags in Sf9-insect cells using a baculovirus/insect cell expression system^31,32^. Pink pellets were washed with PBS, stored at −70 °C, and later purified at 4 °C at all stages. We selected n-Dodecyl-B-D-Maltoside (DDM, Serva Elec.) detergents at around 2x critical micelle concentration (CMC) and supplied Cholesteryl Hemisuccinate (CHS, Serva Elec.) lipids for solubilization of the membrane protein. Cell pellets were resuspended in lysis buffer A, treated with a high-pressure homogeniser (Avestin EmulsiFlex-C3) and samples were centrifuged at 40,000 rpm for one hr. The supernatant was incubated with Ni-NTA beads for four hrs and applied to a polypropylene column (BioRad) equilibrated in lysis buffer A. The column was washed with buffers B and C, and protein was eluted with buffer D. Fractions were isolated based on pink-marker colouration and concentrated using an AMICON ULTRA-15 filter (100 kDa cut-off). The concentrated sample was spun for one hr on a table-top centrifuge at full speed and supernatant was applied onto a Superose 6 Increase 5/150 GL column for gel filtration using buffer E. Buffer compositions are provided in Supplementary Table 2.

**Supplementary Table 2:** Buffers used for protein purification.

| Buffer | Composition |
| --- | --- |
| Lysis buffer A | 50 mM HEPES pH 7.5, 100 mM NaCl, 20 mM imidazole, 1 % w/v DDM, 0.5 % w/v CHS, 1 % protease inhibitor (cOmplete Protease Inhibitor Cocktail EDTA-free, Sigma-Aldrich) |
| Wash buffer B | 50 mM HEPES pH 7.5, 100 mM NaCl, 20 mM imidazole, 0.01 % w/v DDM, 0.005 % CHS, 1 % protease inhibitor |
| Wash buffer C | 50 mM HEPES pH 7.5, 100 mM NaCl, 50 mM imidazole, 0.01 % w/v DDM, 0.005 % CHS, 1 % protease inhibitor |
| Elution buffer D | 50 mM HEPES pH 7.5, 100 mM NaCl, 350 mM imidazole, 0.01 % w/v DDM, 0.005 % CHS, 1 % protease inhibitor |
| Gel filtration buffer E | 50 mM HEPES pH 7.5, 100 mM NaCl, 0.01 % w/v DDM, 0.005 % CHS |

### Mass photometry

Measurements were performed in buffer E (see above) using an One^MP^ mass photometer (Refeyn Ltd, Oxford, UK)^33^. Immediately before analysis, the sample was diluted 1:10 with the aforementioned buffer. Molecular mass was determined in the analysis software provided by the manufacturer using a NativeMark-(Invitrogen).

### Structure modelling

The homology model of the tetrameric Kcnj13 channel (Fig. 5e) was built using SWISS-MODEL^66-70^ based on the crystal structure template (2.6-Å resolution) of the potassium channel Kir2.2 from *Gallus gallus* (PDB ID: 3spg), sharing a sequence similarity of 37 % with the target protein Kcnj13 from *D. rerio*. Similar models with a pTM-based confidence score of ∼ 60 % were generated using AlphaFold-Mutlimer^71,72^.

### Genome and transcriptome sequencing

Reciprocal crosses between species (male *D. aesculapii* x female *D. rerio* (pair 1), and male *D. rerio* x female *D. aesculapii* (pair 2)) were performed via in vitro fertilization to produce F1 hybrids. Adult parental fish (n=4) and F1 hybrids (n=12; 7 hybrids from cross 1, 5 hybrids from cross 2) were euthanized by exposure to buffered 0.5 g/L MS-222 (Tricaine). Tissues were dissected in ice-cold PBS and collected using TRIzol (Life Technologies). DNA from the parental individuals was isolated from posterior trunk tissue including the fins. RNA was obtained from skin and posterior trunk tissue of F1 hybrids. RNA integrity and quantity were assessed by Agilent 2100 Bioanalyzer. Metadata is provided in Supplementary Table 3. Library preparation (DNA/RNA: TruSeq DNA Nano Kit (Illumina); 100 ng per sample) and sequencing (NovaSeq 6000 (Illumina), for DNA: 2x 250 bp, for RNA: 2x 100 bp) were performed by CeGaT GmbH (Tübingen, Germany). Data are available in: PRJEB53585.

All subsequent analyzes were based on high-quality clean reads. Quality of the sequencing data was checked using FastQC (version 0.11.9) and adapter sequences were trimmed using fastp (version 0.23.2)^73^. Genome resequencing reads were aligned to the *D. rerio* reference genome (GRCz11) using BWA-MEM (version 0.7.17-r1188)^74^. The aligned SAM files were sorted and converted into BAM files using SAMtools (version 1.11)^75^. Then the sorted BAM files were de-realigned and indexed again using Picard (version 2.18.29, https://broadinstitute.github.io/picard/). Transcriptomes were aligned to GRCz11 using STAR aligner (version 2.7.10a)^76^. The BAM files directly output by STAR in two-pass mode are deduplicated and indexed by Picard.

### Variant calling and filtration

To identify species-specific alleles, variant calling was performed according to the best practice pipeline of the Genome Analysis Toolkit (GATK4)^77,78^. Specifically, Haplotypecaller was used to detect variants based on genome and transcriptome data. The called variants were joint-genotyped using GentypeGVCFs into a single .vcf file; data from skin and trunk tissue were separately processed. First, SelectVariants was used to filter single nucleotide polymorphisms (SNPs), then the selected SNPs were hard-filtered using Variantfiltration. Specifically, SNPs of ‘QUAL < 30.0, QD < 2, FS > 60, MQ < 40, SOR > 3, MQRankSum < −12.5 and ReadPosRankSum > −8’ as well as non-biallelic SNPs were filtered out. The remaining SNPs were filtered again using VCFtools (--max-missing 0.8, --maf 0.05). Finally, SNPs shared by genomes and transcriptomes were selected for the subsequent allele-specific expression analysis (ASE) using the intersect function of Bedtools (version 2.30.0)^79^.

### Allele-specific expression analysis

Read counts for species-specific SNPs were averaged per gene for each hybrid transcriptome using GATK ASEReadCounter^80^ with default filters enabled. Significant allele-specific expression was defined as ‘Fold Change’ > 2 between alleles and adjusted p-values (p-adj) < 0.05 from DESeq2 package in R^81^. Finally, the ggplot2^82^ package in R rendered a volcano plot using the data obtained by DESeq2.

## Data availability

The authors declare that all data supporting the findings of this study are available within the article and its supplementary information files or from the corresponding author upon reasonable request. The dataset generated during this study is available at The European Nucleotide Archive (ENA) accession number: PRJEB53585.

**Supplementary Table 3:** Metadata for transcriptomic analysis.

| CeGaT ID | short sample description | sex | stage | Egg_lay_date | Sampling_date | Extraction_date | RIN_value | quantity_u<br>g | Concentration_ng_uL |
| --- | --- | --- | --- | --- | --- | --- | --- | --- | --- |
| S1906Nr1 | RNA_skin_rerio-aesculapii_pair1_hybrid_1 | NA | adult | 20190606 | 20200113 | 20190114 | 9,30 | 0,814 | 90,4 |
| S1906Nr2 | RNA_trunk_rerio-aesculapii_pair1_hybrid_1 | NA | adult | 20190606 | 20200113 | 20190114 | 9,30 | 1,752 | 219 |
| S1906Nr3 | RNA_skin_rerio-aesculapii_pair1_hybrid_2 | NA | adult | 20190606 | 20200113 | 20190114 | 8,20 | 1,548 | 172 |
| S1906Nr4 | RNA_trunk_rerio-aesculapii_pair1_hybrid_2 | NA | adult | 20190606 | 20200113 | 20190114 | 8,20 | 1,856 | 232 |
| S1906Nr5 | RNA_skin_rerio-aesculapii_pair1_hybrid_3 | NA | adult | 20190606 | 20200113 | 20190114 | 7,90 | 1,54 | 154 |
| S1906Nr6 | RNA_trunk_rerio-aesculapii_pair1_hybrid_3 | NA | adult | 20190606 | 20200113 | 20190114 | 8,20 | 4,23 | 470 |
| S1906Nr7 | RNA_skin_rerio-aesculapii_pair1_hybrid_4 | NA | adult | 20190606 | 20200113 | 20190114 | 8,50 | 1,43 | 130 |
| S1906Nr8 | RNA_trunk_rerio-aesculapii_pair1_hybrid_4 | NA | adult | 20190606 | 20200113 | 20190114 | 9,30 | 5,7 | 570 |
| S1906Nr9 | RNA_skin_rerio-aesculapii_pair1_hybrid_5 | NA | adult | 20190606 | 20200113 | 20190114 | 8,20 | 1,6 | 160 |
| S1906Nr10 | RNA_trunk_rerio-aesculapii_pair1_hybrid_5 | NA | adult | 20190606 | 20200113 | 20190114 | 7,80 | 3,652 | 332 |
| S1906Nr11 | RNA_skin_rerio-aesculapii_pair1_hybrid_6 | NA | adult | 20190606 | 20200113 | 20190114 | 8,50 | 1,118 | 93,2 |
| S1906Nr12 | RNA_trunk_rerio-aesculapii_pair1_hybrid_6 | NA | adult | 20190606 | 20200113 | 20190114 | 9,20 | 3,204 | 356 |
| S1906Nr13 | RNA_skin_rerio-aesculapii_pair1_hybrid_7 | NA | adult | 20190606 | 20200113 | 20190114 | 9,00 | 1,232 | 112 |
| S1906Nr14 | RNA_trunk_rerio-aesculapii_pair1_hybrid_7 | NA | adult | 20190606 | 20200113 | 20190114 | 9,60 | 1,76 | 160 |
| S1906Nr15 | RNA_skin_rerio-aesculapii_pair2_hybrid_8 | NA | adult | 20190814 | 20200113 | 20190114 | 9,20 | 1,56 | 120 |
| S1906Nr16 | RNA_trunk_rerio-aesculapii_pair2_hybrid_8 | NA | adult | 20190814 | 20200113 | 20190114 | 9,40 | 3,77 | 290 |
| S1906Nr17 | RNA_skin_rerio-aesculapii_pair2_hybrid_9 | NA | adult | 20190814 | 20200113 | 20190114 | 7,70 | 1,596 | 114 |
| S1906Nr18 | RNA_trunk_rerio-aesculapii_pair2_hybrid_9 | NA | adult | 20190814 | 20200113 | 20190114 | 9,90 | 2,288 | 176 |
| S1906Nr21 | RNA_skin_rerio-aesculapii_pair2_hybrid_11 | NA | adult | 20190814 | 20200113 | 20190114 | 8,80 | 1,644 | 137 |
| S1906Nr22 | RNA_trunk_rerio-aesculapii_pair2_hybrid_11 | NA | adult | 20190814 | 20200113 | 20190114 | 9,70 | 4,296 | 358 |
| S1906Nr25 | RNA_skin_rerio-aesculapii_pair2_hybrid_13 | NA | adult | 20190814 | 20200113 | 20190114 | 9,40 | 1,365 | 105 |
| S1906Nr26 | RNA_trunk_rerio-aesculapii_pair2_hybrid_13 | NA | adult | 20190814 | 20200113 | 20190114 | 10,00 | 1,728 | 144 |
| S1906Nr27 | RNA_skin_rerio-aesculapii_pair2_hybrid_14 | NA | adult | 20190814 | 20200113 | 20190114 | 9,50 | 1,508 | 116 |
| S1906Nr28 | RNA_trunk_rerio-aesculapii_pair2_hybrid_14 | NA | adult | 20190814 | 20200113 | 20190114 | 9,40 | 3,406 | 262 |
| S1906Nr29 | DNA_D_aesculapii_parent3_male_pair1 | male | adult | NA | 20190612 | 20190612 | NA | 2,104 | 11,5 |
| S1906Nr30 | DNA_D_rerio_parent1_female_pair1 | female | adult | NA | 20190612 | 20190612 | NA | 5,746 | 31,4 |
| S1906Nr31 | DNA_D_rerio_parent5_male_pair2 | male | adult | NA | 20190816 | 20190816 | NA | 7,997 | 43,7 |
| S1906Nr32 | DNA_D_aesculapii_parent9_female_pair2 | female | adult | NA | 20190816 | 20190816 | NA | 5,124 | 28 |

## Acknowledgements

We thank Hans-Martin Maischein (now Max Planck Institute for Heart and Lung Research, Bad Nauheim, Germany) and Horst Geiger (Max Planck Institute for Biology, Tübingen, Germany) for help with blastula transplantations, Christian Feldhaus and Aurora Panzera (BioOptics Facility, Max Planck Institute for Biology) for help with imaging of the transgenic lines, Veronika Altmannova and Dorota Rousova (Friedrich Miescher Laboratory, Tübingen, Germany) for help with protein purification, and Silke Geiger-Rudolph, Roberta Occhinegro and Reinhard Albrecht for excellent technical assistance (Max Planck Institute for Biology, Tübingen, Germany). The AlphaFold-Multimer model was generated using the BMBF-funded de.NBI Cloud within the German Network for Bioinformatics Infrastructure (de.NBI) (031A532B, 031A533A, 031A533B, 031A534A, 031A535A, 031A537A, 031A537B, 031A537C, 031A537D, 031A538A). This work was supported by an ERC Advanced Grant “DanioPattern” (694289) and the Max Planck Society, Germany.

## Contributions

M.P., A.P.S., C.M.D., H.G.F., S.W., C.N.V. and U.I. were involved in the design of the experiments. M.P., A.P.S., U.I., H.G.F., and M.F. performed the experiments. U.I., M.P., C.N.V., A.P.S., J.L., Z.F., C.M.D., H.E., S.W., J.R.W. and analysed the data. M.P. made the figures with help from U.I. and C.N.V.; M.P., U.I., A.P.S. and C.N.V. wrote the manuscript. C.N.V. and J.R.W. acquired funding.

